# Early transcriptional similarities between two distinct neural lineages during ascidian embryogenesis

**DOI:** 10.1101/2023.12.20.572557

**Authors:** Richard R Copley, Julia Buttin, Marie-Jeanne Arguel, Géraldine Williaume, Kevin Lebrigand, Pascal Barbry, Clare Hudson, Hitoyoshi Yasuo

## Abstract

In chordates, the central nervous system arises from precursors that have distinct developmental and transcriptional trajectories. Anterior nervous systems are ontogenically associated with ectodermal lineages while posterior nervous systems are associated with mesoderm. Taking advantage of the well-documented cell lineage of ascidian embryos, we asked how the transcriptional states of the different neural lineages become similar during the course of progressive lineage restriction. We performed single-cell RNA sequencing (scRNA-seq) analyses on hand-dissected neural precursor cells of the two distinct lineages, together with those of their sister cell lineages, with a high temporal resolution covering five successive cell cycles from the 16-cell to neural plate stages. A transcription factor binding site enrichment analysis of neural specific genes at the neural plate stage revealed limited evidence for shared transcriptional control between the two neural lineages, consistent with their different ontogenies. Nevertheless, PCA analysis and hierarchical clustering showed that, by neural plate stages, the two neural lineages cluster together. Consistent with this, we identified a set of genes enriched in both neural lineages at the neural plate stage, including *miR-124, CELF3/5/6, Zic.r-b,* and *Ets1/2*.

## Introduction

The dorsal neural tube of the central nervous system (CNS) is a synapomorphy of chordates (Satoh et al., 2014). In both vertebrate and invertebrate chordates, neural cells of anterior and posterior CNS have followed distinct developmental and transcriptional trajectories (Gouti et al., 2015, 2014; Henrique et al., 2015; Hudson and Yasuo, 2021). Using ascidian embryos, we wanted to address the extent that the transcriptional states of two distinct neural lineages, arising from distinct embryonic origins, become similar during early chordate embryogenesis. Ascidians are invertebrate chordates that develop a well-patterned dorsal CNS at larval stages. This consists of a sensory vesicle, or brain, followed by a trunk ganglion and tail nerve cord (Fig 1A). The structure and underlying specification mechanisms of the ascidian larval CNS are well documented (reviewed in (Hudson, 2016; Hudson and Yasuo, 2021; Liu and Satou, 2020; Ryan and Meinertzhagen, 2019)). Uniquely among chordates, ascidian embryos develop with an invariant cleavage pattern and their cell lineages are well described. At the 8-cell stage of development, the four founder lineages arise with a-and b-line cells in the animal half, contributing to predominantly ectodermal lineages, and A-and B-line cells in the vegetal half, contributing to predominantly mesendodermal lineages (Nishida, 1987). The anterior part of the sensory vesicle (=brain) originates from the a-line ectodermal lineage following a cell fate choice between neural and epidermal lineages. The dorsal-most row of cells, from the posterior part of the sensory vesicle to the tail nerve cord, derives from the b-line, following fate choices between neural and epidermal or neural and muscle lineages. The remainder of the CNS, all lateral and ventral cells from the posterior part of the sensory vesicle to the tail nerve cord, originates from the A-line mesendodermal lineage following fate choices between neural and notochord or neural and muscle lineages. While the a-and A-line neural lineages originate from the animal and vegetal hemispheres of the embryo respectively, they are found juxtaposed at the “dorsal” marginal zone of early embryos and then collectively form the neural plate (Fig 1B).

**Fig 1.**
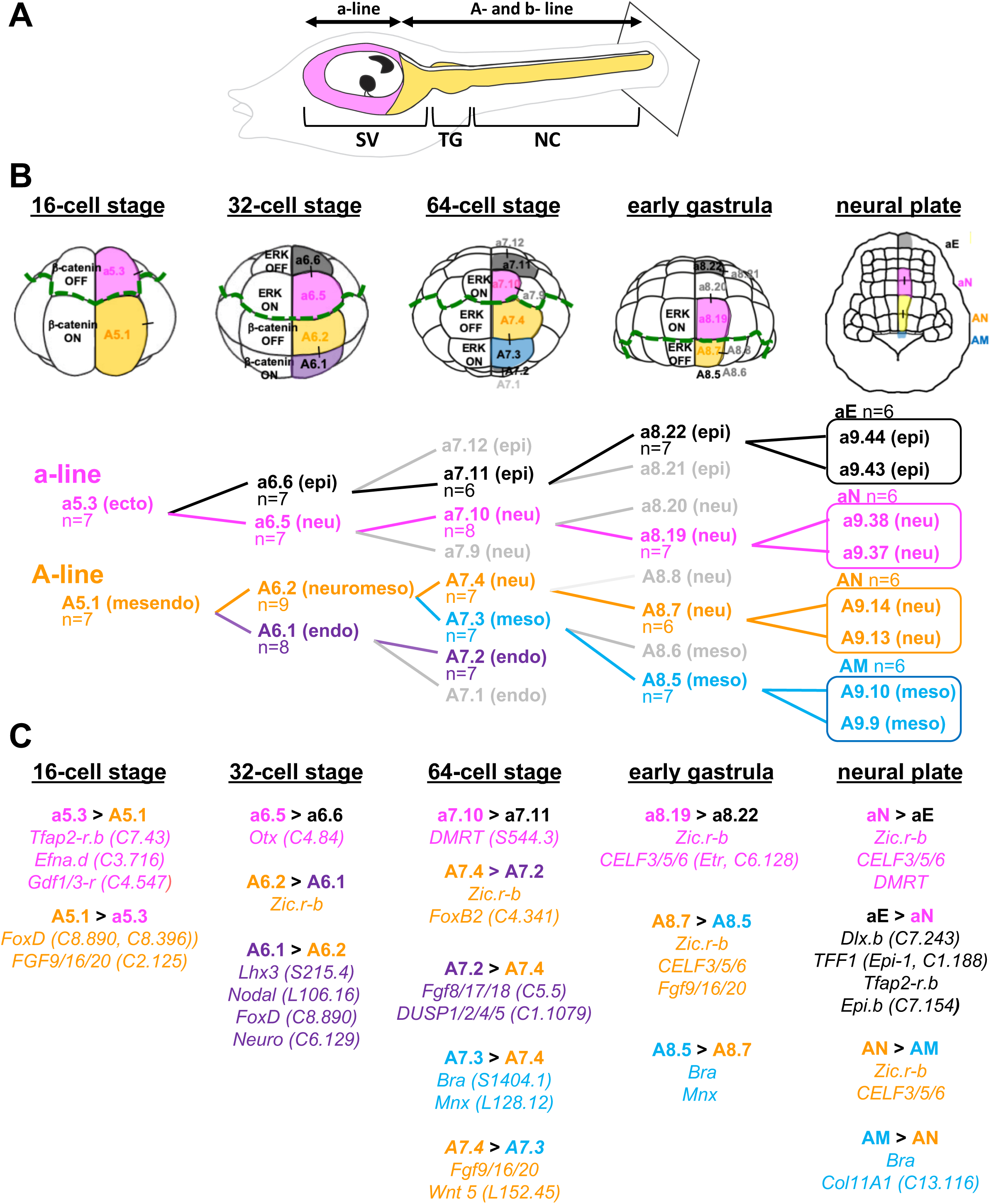
Two distinct developmental lineages for *Ciona* larval CNS: a-line and A-line. (A) Drawing of the larval CNS enlarged within the larvae outline. The tail is truncated (indicated by the square). The larval CNS is derived from a-line cells (pink), which make up the anterior part of the sensory vesicle and A-line cells (orange) which make up much of the remaining CNS. The dorsal most row of cells from the posterior sensory vesicle to the tip of the tail arises from the b-lineage (not coloured). SV: sensory vesicle; TG: trunk ganglion; NC: nerve cord. (B) Isolated cells from each stage were coloured with a-and A-line neural lineages in pink and orange, epidermis in dark grey, mesoderm in light blue and endoderm in purple. Black lines connecting two given cells indicate their sister relationship. Note that cells of the neural plate stage embryo were isolated as pairs of two sister cells. n= number of each particular cell type sequenced. (C) The presence of expected differentially expressed genes (“cell 1>cell 2” indicates a set of genes enriched in cell 1) confirmed correct identification and isolation of the cells. *Zic.r-b* is a multi-copy gene (KH.S816.1, KH.S816.2, KH.S816.4, KH.L59.1, and KH.L59.12).

Segregation of neural fate within the different lineages depends upon a distinct sequence of signalling inputs. Neural induction in the a-line, mediated by FGF signals, begins at the 32-cell stage with transcriptional induction of *Otx* in a6.5 cells, segregating the neural lineage (CNS and placode) from ectoderm (Bertrand et al., 2003; Nishida, 1987). a-line neural precursors become CNS-specific at the early gastrula stage when the CNS-neural precursors segregate from the precursors of a specialized proto-placodal region of the anterior ectoderm (Abitua et al., 2015; Wagner and Levine, 2012). From the 16-cell stage of development, the a-line neural cells are exposed to a series of signalling inputs (Fig 1B). At the 16-cell stage, they are found in a territory in which the canonical β-catenin signalling pathway is inactive (β-catenin OFF) (Imai et al., 2000; Rothbacher et al., 2007). From the 32-cell to early gastrula stage, the ERK-signalling pathway is active (ERK-ON) in the a-line neural precursors (Hudson et al., 2003; Nishida, 2003; Wagner and Levine, 2012). The A-line neural lineages segregate at the 64-cell stage from the sister mesodermal lineage and follow a distinct sequence of signalling inputs compared to their a-line counterparts. At the 16-cell stage, A-line neural precursors are found in the β-catenin-ON territory, followed by β-catenin-OFF and ERK-ON at the 32-cell stage and then ERK-OFF from the 64-cell to early gastrula stage (Hudson et al., 2003; Imai et al., 2000; Minokawa et al., 2001; Picco et al., 2007; Rothbacher et al., 2007). These distinct sequences of combinatorial signalling inputs are required for the correct specification of neural precursors from the two neural lineages and the ON-OFF status of these signal inputs dictates their binary cell fate decisions. Thus, a-line and A-line neural precursors derive from different lineages and are specified by distinct molecular mechanisms. Across the medial-lateral axis of the neural plate, both neural lineages are patterned by Nodal and Delta signals (Esposito et al., 2016; Hudson et al., 2007). In this manuscript, we focused on the medial neural precursors that do not depend on Nodal signals (Fig 1B).

Single-cell RNA sequencing (scRNA-seq) analyses have been used in ascidians to study the transcriptome trajectories of differentiating embryonic cells and mostly recapitulate known developmental lineage segregations (Cao et al., 2019; Sladitschek et al., 2020; Winkley et al., 2021; Zhang et al., 2020). In particular, during development of the CNS lineages from the early gastrula to larva stages, a progressive increase in cell type complexity was revealed, with 41 neural subtypes identified at larval stages (Cao et al., 2019). A trajectory inference analysis was able to assign each of these neural subtypes to a specific developmental lineage origin, indicating that differentiating neural precursors retain transcriptional “identities” that can be connected, through a sequence of intermediates, to their developmental origins (Cao et al., 2019). These previous scRNA-seq studies have highlighted the diversification of cell types during ascidian embryogenesis with single cell transcriptome trajectories following developmental lineage segregations.

In this study, by applying the scRNA-seq approach to early developmental stages of ascidian embryogenesis, we addressed whether the transcriptional states of the a-and A-line neural lineages become similar to each other while diverging from their sister lineages. Following the medial a-and A-line cells with a high temporal resolution of each cell cycle from the 16-cell to the neural plate (mid-gastrula) stages, using scRNA-seq of hand-dissected cells, we looked for evidence of a shared “pan-neural” transcriptional state. While neural lineage cells cluster with their corresponding sister lineage cells up to early gastrula stage, we found that, by mid-gastrula (neural plate stage), the a-and A-line neural cells cluster together, indicating that these distinct neural lineages are converging in some aspects of their transcriptional states. Consistent with this observation, we could identify a set of genes significantly enriched in both neural lineages. Further, we could not detect lineage-based clustering between the 41 larval neural cell type endpoints (Cao et al., 2019), supporting the notion that during development of the CNS, the transcriptional state of the cells becomes dominated by the functional end point rather than the lineage origin.

## Results and Discussion

### Isolation of neural and sister cell lineages from the 16-cell stage to neural plate stage of development

We used *Ciona robusta* (*C. intestinalis type A*) since its genome is better annotated than that of *Ciona intestinalis* (*C. intestinalis type B*) (Satou et al., 2021, 2019). In order to generate scRNA-seq datasets for individual neural lineage cells of each cell cycle from the 16-cell to neural plate stages, we manually dissected cells of interest. In parallel, we also isolated cells of the corresponding early segregating sister lineages. All the isolated cells are shown in Figure 1B (Fig 1B). While hand-dissection has the disadvantage that it is only feasible to isolate limited numbers of cells, a clear advantage of the approach is that the precise identity of the cell is known and that cells can be frozen immediately following isolation. Neural lineage precursors, a5.3 and A5.1, were dissected at the 16-cell stage. At the 32-cell stage, we isolated neural a6.5 and epidermal a6.6 sister precursors from the a-line and neuromesodermal A6.2 and endodermal A6.1 sister cells from the A-line (Fig 1B). At the 64-cell stage, the neural precursor a6.5 has divided in a medio-lateral direction and we isolated the medial neural precursor a7.10 and the epidermal precursor a7.11, a daughter cell of a6.6. In the A-line, the neuromesodermal cell has divided asymmetrically, generating a notochord precursor A7.3 and a neural precursor A7.4. We isolated each of these sister cells, together with the A7.2 endoderm precursor, a daughter cell of A6.1. At the early gastrula stage corresponding to the 112-cell stage, the a7.10 neural precursor has divided into a CNS precursor a8.19 and a placodal precursor a8.20. We isolated the a8.19 neural cell together with a8.22 epidermal precursor, which is a daughter cell of a7.11. In terms of the A-line of the early gastrula, both neural and notochord precursors have divided medio-laterally by this stage and we isolated the medially positioned neural precursor A8.7 and notochord precursor A8.5. At the 6-row neural plate stage, the neural plate exhibits a grid-like structure with six rows of cells along the A-P axis (Fig 1B). This results from all neural precursors of the early gastrula stage dividing along the anterior-posterior axis. At this stage, differential ERK activation takes place between row I and row II cells of the A-line neural precursors and between row III and row IV of the a-line neural precursors (Haupaix et al., 2014; Nishida, 2003). Since we wanted to compare the a-line and A-line neural lineages globally, and for technical facility, we isolated medial neural plate cells as pairs, row I/row II (A-line) and row III/row IV (a-line), which we called AN (neural, daughters of A8.7) and aN (neural, daughters of a8.19), and their counterpart sister lineage cells, AM (notochord, daughters of A8.5) and aE (epidermis, daughters of a8.22) (Fig 1B). The total number of manually isolated cells (or cell pairs for aN, aE, AN, and AM) resulted in a 6-9-fold coverage of each cell type (Fig 1B). Altogether, the sampled cells represent five successive cell cycles of the a-and A-neural lineages and of their sister lineages, covering the 16-cell to the neural plate stages.

### Differential enrichment of transcriptomes between pairs of cells

As a consequence of the hand dissection procedure, our cell transcriptomes were already labelled by cell type of origin. All our downstream computational analyses were based on the KH2013 gene models (Satou et al., 2008). We analysed cells for differential expression between neural and sister lineages using DESeq2, treating each cell (n = 6 to 9) as a single replicate instance of its type (see supplementary data link at end of manuscript). Inspection of principal component analysis (PCA) plots of rlog-transformed gene expression levels for cells of a particular embryonic stage revealed a strong tendency of cells isolated at the early stages (16-and 32-cell stages) to cluster by batch or embryo of origin, as previously observed by others in *Ciona* (Ilsley et al., 2020; Winkley et al., 2021). Accordingly, for these stages, we added the animal of origin to the DESeq2 design formula. Genes were ranked by adjusted P-values and values < 0.01 regarded as significant (S1 Table: DESeq2). Our DESeq2 analysis successfully recovered genes whose expressions were previously reported to be differentially expressed between the pairs of cells analysed in the current study (Fig 1C), showing that manual dissections were conducted with accurate cell identification.

### Trajectory of transcriptional states during the course of neural lineage segregation up to neural plate stages

A PCA of the variance stabilized (vst) expression data for all cells showed two dominant themes: 1) PC 1 reflecting embryonic stage or time and 2) PC 2 reflecting the different lineages (Fig 2A). Separation between lineages becomes stronger through developmental time. At the neural plate stage, separation is strongest between the aE and AM lineages, with the neural aN and AN lineages appearing relatively closer to each other. To investigate this effect further, we performed a hierarchical clustering analysis (see methods and Fig 2B). This showed a clustering of the aN and AN cells of the neural plate stage as each other’s closest neighbours, in contrast to neural precursors at earlier stages that clustered with their sister lineage cells. These data suggest that, as lineage segregation proceeds, transcriptional similarity attributable to shared lineage history is overwritten by similarity caused by the acquired developmental identity of the cell. In order to further address this converging trend, we used the available scRNA-seq dataset of larval neural cell types, which emerge following a maximum of four rounds of cell divisions of neural plate cells (Cao et al., 2019), and clustered them based on the shared presence or absence of marker genes. The hierarchical clustering showed no obvious segregation of these CNS neural cell types by lineage of origin (‘A’ or ‘a’) (Fig 3).

**Fig 2.**
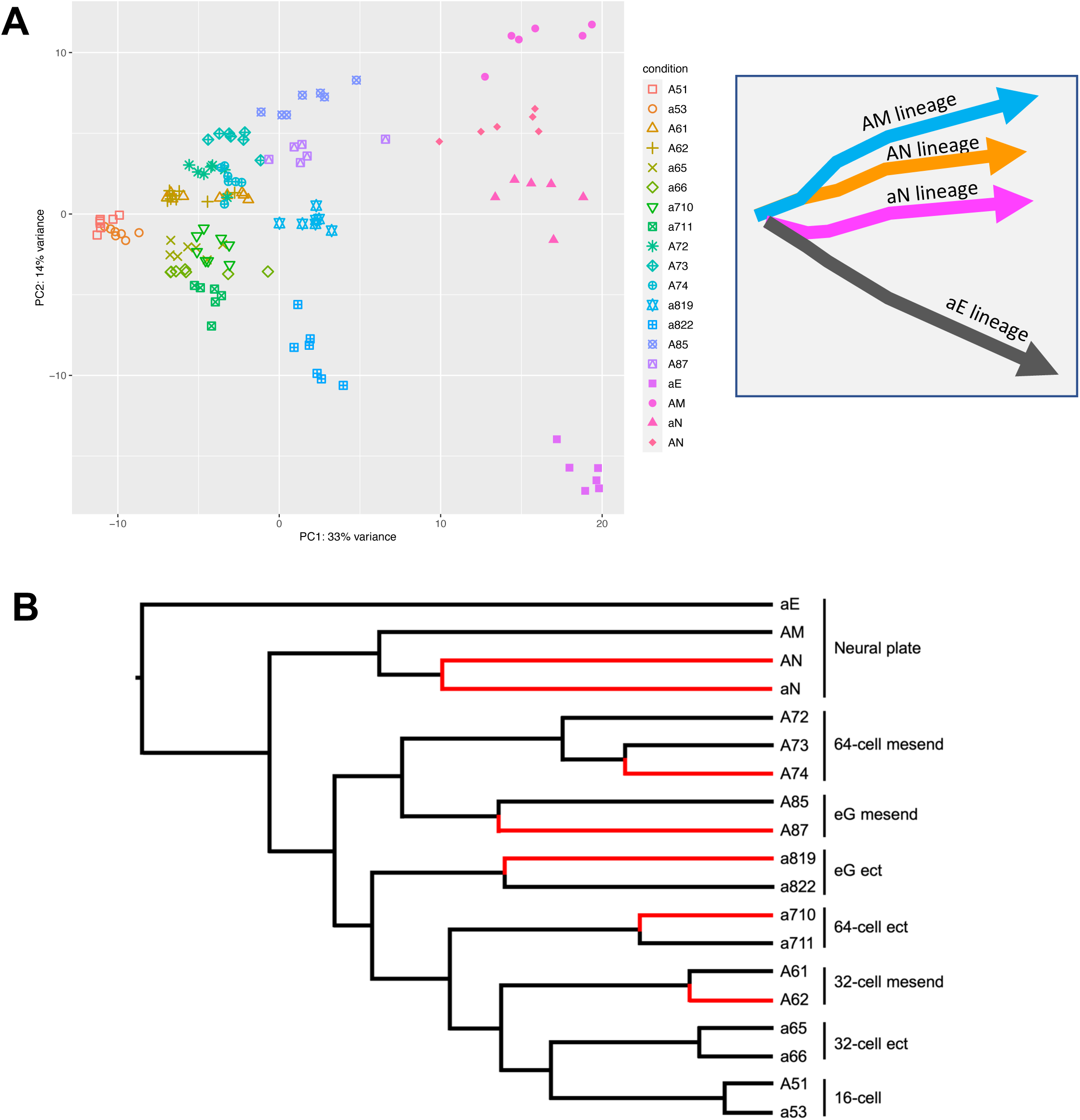
Transcriptional trajectories during the course of neural lineage segregation. (A) Principal component analysis (PCA) of the cells (left). Count data per gene were variance stabilizing transformed (vst in DESeq2, see methods) and the 500 genes with greatest variance used for the basis of the PCA. Each point represents a single cell following the key on the right side that indicates its identity. Schematic drawing (right) represents the transcriptional trajectories of the different lineages analysed. (B) Cluster tree of the cell types (see methods for details). With the exception of aN and AN, the closest neighbour of all neural lineage precursors is its corresponding early diverging sister lineage cell. Note that, at the 16-cell stage, the a5.3 and A5.1 cells have no sampled sister cell types of the same lineage. Neural lineages are represented by red branches. eG: early gastrula; ect: ectoderm; mesend; mesoendoderm (cf. Fig 1).

**Fig 3.**
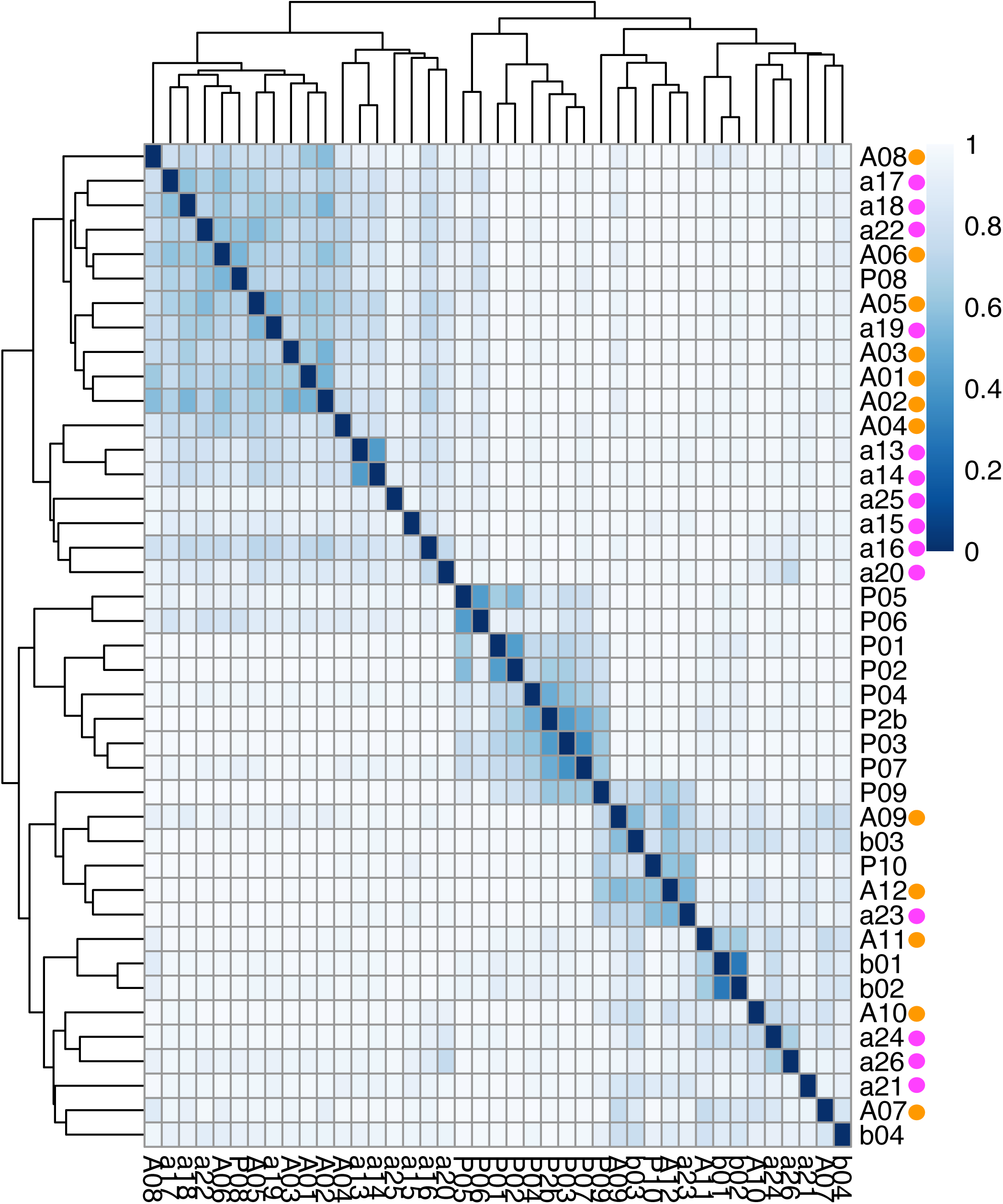
Hierarchical clustering of larval stage neural cell types. In order to highlight their respective lineage origins, the larval neural cell clusters defined in Cao et al, 2019 are renamed as follows (“A” for A-line neural lineage, “a” for a-line neural lineage, “b” for b-line neural lineage, and “P” for peripheral nervous system): A01 *KCNB1*+ motor ganglion; A02 *GLRA1*+ motor ganglion; A03 *AMD*+ motor ganglion; A04 *VP*+ pSV; A05 *VP-R*+ SV; A06 *GSTM1*+ SV; A07 MHB; A08 Tail nerve cord (A); A09 Ependymal cells; A10 *GLGB*+ pSV; A11 Trunk nerve cord (A); A12 *Pax2/5/8-A*+ neck; a13 *Arx*+ pro-aSV; a14 *Aristaless*+ aSV; a15 *Opsin1*+, *PTPRB*+ aSV; a16 *Rx*+ aSV; a17 *FoxP*+ aSV; a18 *Opsin1*+, *STUM*+ aSV; a19 *Lhx1*+ GABAergic neurons; a20 *Lox5*+ aSV; a21 *Lhx1*+, *Bsh*+ aSV; a22 Eminens; a23 *Six3/6*+ pro-aSV; a24 *Hedgehog2*+ SV; a25 Coronet cells; a26 Pigment cells; b01 Tail nerve cord (b); b02 *Arx*+ nerve cord (b); b03 Trunk nerve cord (b); b04 *FoxD-b*+ cells; P01 Glia cells; P02 pATENs; P2b PSCs related; P03 Collocytes; P04 PSCs; P05 CESNs; P06 RTENs; P07 aATENs; P08 BTNs; P09 *Dll-A*+ ANB (anterior neural boundary); P10 *Pitx*+ ANB (anterior neural boundary). Furthermore, neural cell clusters of the A-line origin are marked with orange dots while those of the a-line origin with pink dots. The transcriptome dataset used in this analysis was generated in Cao et al, 2019.

Altogether, we observe that the transcriptional states of the two neural lineages become more similar to each other than they are to their early diverging sister lineages by the neural plate stage of development (Fig 2) followed by extensive mixing of differentiated neural cell types derived from these distinct lineages at larval stages (Fig 3). In other words, the transcriptional signature of their lineage identity becomes weaker as development proceeds.

### Identification of neural genes whose transcripts are enriched in both neural lineages

The above data suggest that neural cells from both lineages shared some aspects of their transcriptional state at the neural plate stage (Fig 2). We therefore searched for transcripts enriched in both neural lineages in our DESeq2 analysis (S1 and S2 Tables). We identified six genes whose transcripts are enriched in both of the neural lineages at the early gastrula stage and 11 genes at the neural plate stages. (S2 Table: shared neural genes) (Figs 4 and 5). This list contains many genes already known to be expressed in developing neural tissue: *CELF3/5/6* (KH.C6.128, an ELAV family RNA-binding protein), *Zic.r-b,c,d,e,f* (multicopy gene: KH.S816.1, KH.S816.2, KH.S816.4, KH.L59.1, and KH.L59.12, a transcription factor), *Ets1/2* (KH.C10.113, a transcription factor), *Noggin* (KH.C12.562, a secreted signalling molecule and antagonist of BMP), and *Pans/miR-124* (KH.C7.140, see later) (Alfano et al., 2007; Brozovic et al., 2018; Chen et al., 2011; Fujiwara et al., 2002; Gainous et al., 2015; Hudson et al., 2003; Hudson and Yasuo, 2005; Imai et al., 2004, 2002; Mita and Fujiwara, 2007). However, the list also includes genes whose expression in neural lineages at the neural plate stage has not been reported. We confirmed expression of some of these genes, with most of them detected in both a-and A-line neural cells (Fig 5). *Noggin* was previously reported to be expressed broadly in the CNS at tailbud stages (Imai et al., 2004). At neural plate stages, we found that *Noggin* is expressed at higher levels in the a-line neural precursors compared to A-line neural plate precursors (Figs 4 and 5). *SLC35F* (KH.C4.90), encoding a putative solute carrier family 35, and *ZCCHC24* (KH.C14.310), encoding a CCHC-type zinc finger protein, were detected in neural plate cells of both a-and A-line lineages at the neural plate stage (Fig 5). Expression of *FAM167* (KH.C14.33), encoding a protein of unknown function, and *PQLC2* (KH.L18.113), encoding a lysosomal cationic amino acid transporter, were detected in both a-and A-line neural precursors at the early gastrula and neural plate stages (Fig 5). *CELF3/5/6*, *Noggin* and *ZCCHC24* are also reported to be expressed in developing central nervous system of vertebrates (Gallo and Spickett, 2010; Kang et al., 2012; Knecht et al., 1995; McMahon et al., 1998).

**Fig 4.**
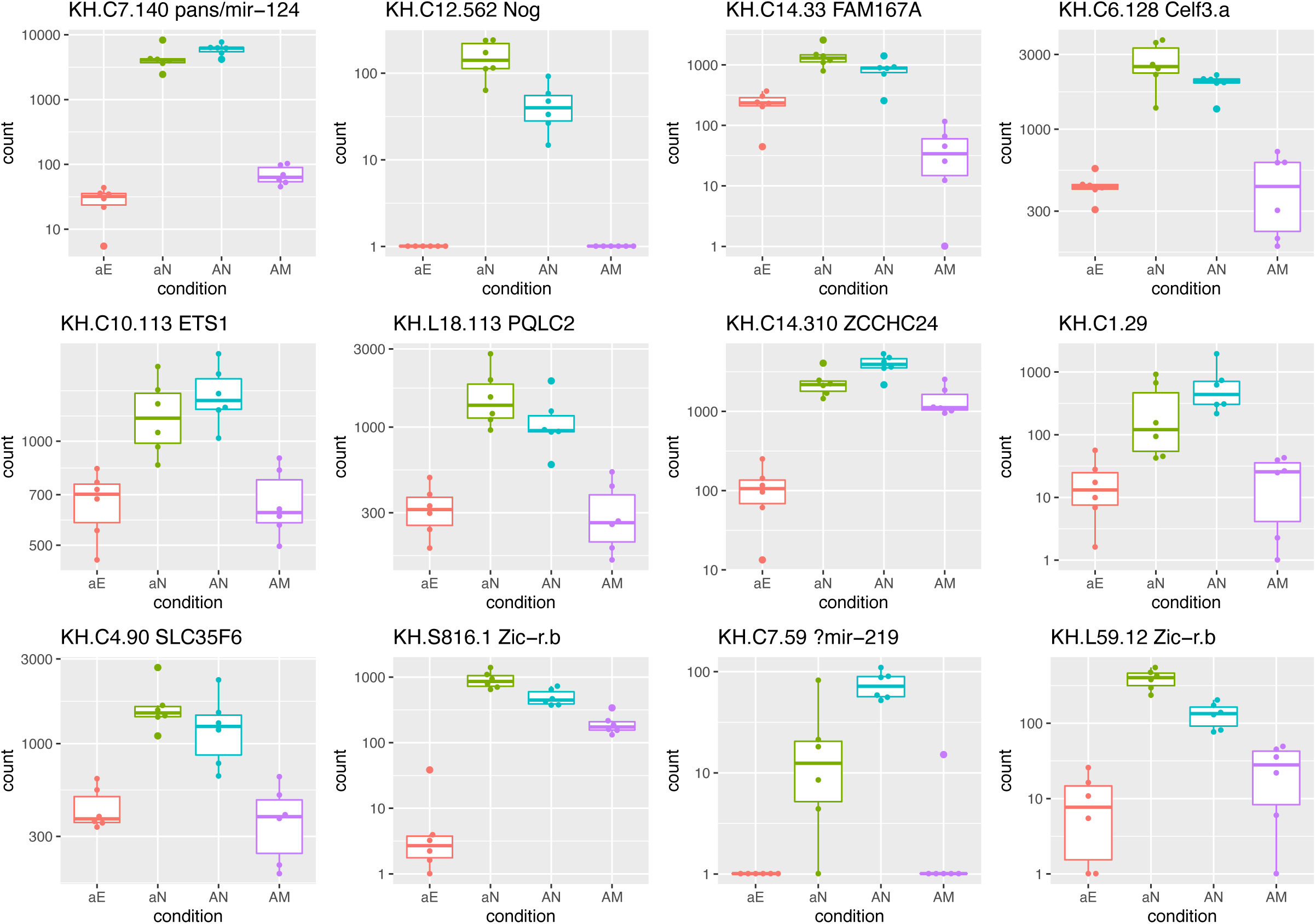
Genes whose transcripts are enriched in both neural lineages at neural plate stage. Graphs show normalized transcript counts in the a-line neural (pink), a-line epidermis (green), A-line neural (blue) or A-line mesoderm (purple) at the neural plate stage. In all cases the comparisons between aN and aE and between AN and AM show statistically significant up regulation (see text for details). Note logarithmic y-axes. Individual data points are represented as dots.

**Fig 5.**
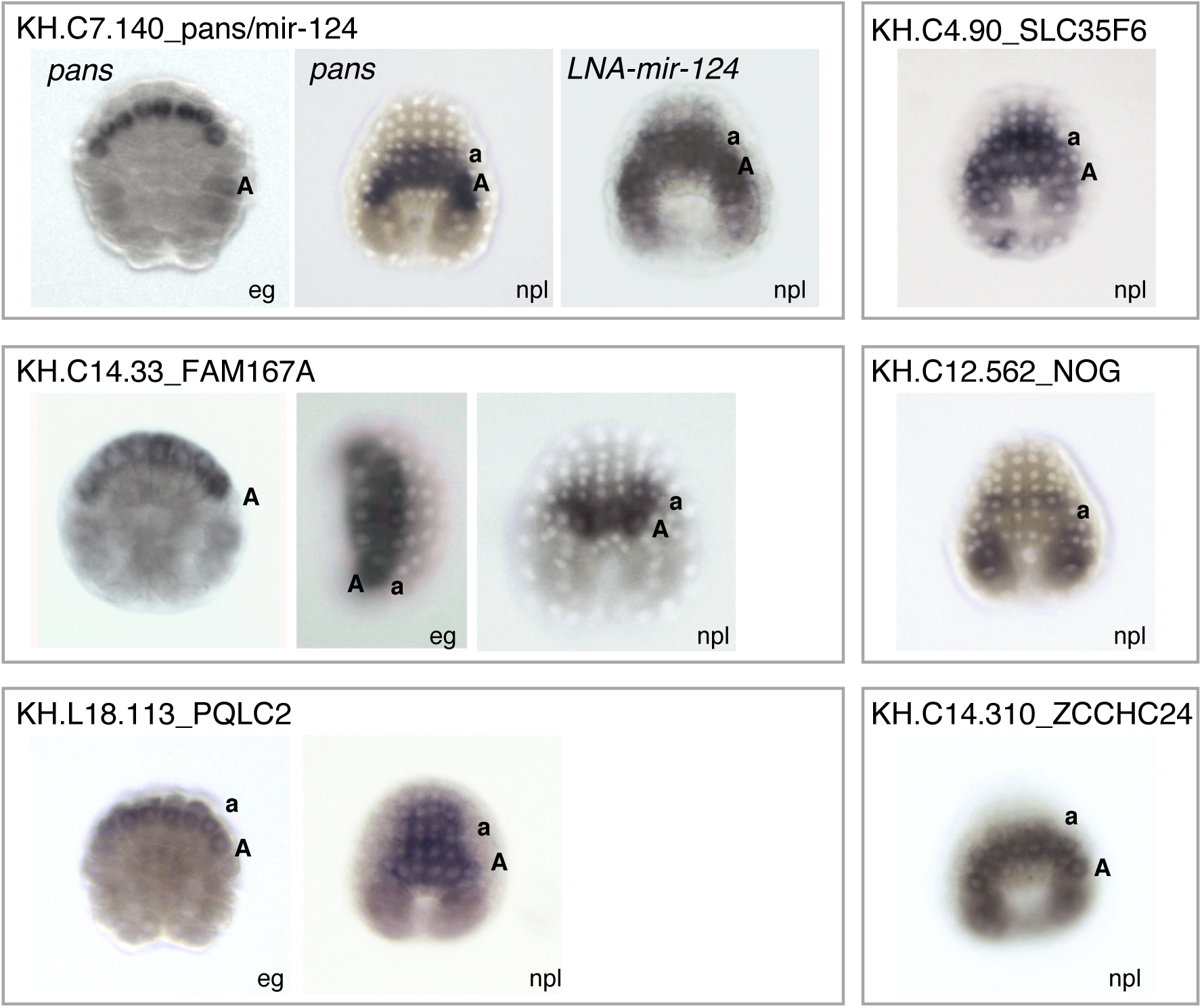
Spatial expression pattern of selected neural-enriched genes at early gastrula and neural plate stages. For all genes analysed, except for *Noggin (NOG)*, expression in both neural lineages was confirmed. A= expression detected in A-line neural cells; a= expression detected in a-line neural cells; eg= early gastrula stage; npl= 6-row neural plate stage.

For the gene encoding *Pans/miR-124* (KH.C7.140), *Pans* expression has been reported in the neural lineages of *Ciona* from the 64-cell stage until tailbud stages (Alfano et al., 2007; Chen et al., 2011; Fujiwara et al., 2002). The predicted *Pans* protein coding gene is very short (22 amino acids) and a role in neural specification has not been reported (Alfano et al., 2007). Within the second intron of this gene, are two tandem copies of the *miR-124* micro-RNA (Chen et al., 2011). The *miR-124* family is highly conserved across metazoans with its expression enriched in nervous systems, in particular, in neuronal lineages (Aboobaker et al., 2005; Clark et al., 2010; Konrad and Song, 2023; Lagos-Quintana et al., 2002; Rajasethupathy et al., 2009; Vidyanand et al., 2017). In *Ciona* embryos, previously published *in situ* hybridisation against the mature *miR-124* product revealed an expression pattern very similar to *Pans* itself (Chen et al., 2011). We observed pan-neural expression of the mature *miR-124* product at the neural plate stage (Fig 5). In vertebrates, *miR-124* expression coincides with neurogenesis, being largely restricted to committed neuronal precursors and post-mitotic neurons, and may play a role in accelerating neuronal differentiation (Visvanathan et al., 2007). In ascidians, *miR-124* appears to be involved in the specification of peripheral epidermal sensory neurons (ESNs) in the epidermal midline: misexpression of *miR-124* in epidermal lineages results in formation of extra ESNs (Chen et al., 2011). It has also been suggested to play a role in downregulation of mesoderm genes, such as *macho-1* (*Zic-r.a*) and notochord genes, to prevent expression of these genes in the CNS lineages (Chen et al., 2011). To further investigate the potential role of *miR-124* in neural specification, we searched for *miR-124* seed sites in the 3’UTRs of genes identified as differentially expressed at the neural plate stage (see methods). In the a-line comparisons, we found an enrichment of *miR-124* binding sites in genes that were upregulated in epidermal lineage cells (aE>aN, p=0.008; aN>aE, p≈1.0). In the A-line comparisons, we found an enrichment of *miR-124* binding sites in genes that were upregulated in mesodermal cells (AM>AN, p=0.0001; AN>AM, p=0.4). It is possible then that *miR-124* might be “fine-tuning” neural lineage segregation by supressing sister-lineage genes as has been observed for other *miRs,* though this will require additional studies (Alberti and Cochella, 2017).

KH.C1.29 encodes a translation initiation factor EIF4EBP1/3. The predicted transcript of KH.C7.59 is very small (312bp; 18 amino acids) and in close proximity to *miR-219* (Brozovic et al., 2018; Hendrix et al., 2010) (S2 Table). *miR-219* has been implicated in some aspects of neural development in vertebrates (Dugas et al., 2010; Hudish et al., 2013).

Among this list of genes enriched in both neural lineages, there are only two genes encoding transcription factors, namely, *Ets1/2* and *Zic.r-b*. Ets1/2 is known to be directly phosphorylated by MAP kinase ERK1/2, leading to an increase in its transcriptional activity, and roles for ERK signalling and Ets1/2 have been described during ascidian nervous system development (Bertrand et al., 2003; Gainous et al., 2015; Haupaix et al., 2014; Ikeda et al., 2013; Racioppi et al., 2014; Yang et al., 1996). *Zic.r-b* has also previously been shown to play a critical role in neural specification (Imai et al., 2006, 2002).

### Transcription factor binding site enrichment in neural lineages at the neural plate stage

The distinct mechanisms of neural lineage segregation in the a-and A-line suggests that neural gene programs would be induced under the influence of different gene regulatory networks. However, the enrichment of expression of two transcription factors, *Zic.r-b* and *Ets1/2* in both neural lineages, described above, suggests that there may also be some common mechanisms by the neural plate stage. In order to investigate this further, we looked for enrichment of transcription factor binding site motifs in the 1kb upstream sequences of differentially expressed genes at neural plate stages (Fig 6 and S3 Table). Each of the four pairwise comparisons, that is, aN>aE, aE>aN, AN>AM, and AM>AN, revealed a unique signature of enriched motifs. Notably, a significant enrichment of Otx-binding sites was found in the genes of both neural lineage comparisons (aN>aE and AN>AM but not aE>aN and AM>AN). Our DESeq2 analyses support differential expression of the *Otx* gene (KH.C4.84) in AN>AM comparison but not in aN>aE comparison (S1 Table: DESeq2). Inspection of the underlying read counts for *Otx* suggests that it is expressed at higher levels in aN>aE but these counts show higher variance, such that the aN>aE comparison is not significant under the DESeq2 statistical model (S1 Fig). *In situ* hybridisation with *Otx* probes shows strong expression in a-line neural lineages from the 32-cell stage, and in both neural lineages from neurula stage (Hudson and Lemaire, 2001; Hudson et al, 2003). In contrast to the neural lineage specific enrichment of Otx-binding sites, the epidermal lineage (aE) showed very strong enrichment for AP2-binding sites. Consistent with this, *Tfap2-r.b* (KH.C7.43), one of the two *Ciona AP2-like* genes, has been shown to play a key role in epidermal differentiation in ascidian embryos (Imai et al., 2016, 2004) and its transcripts are enriched in epidermal lineages (aE>aN) (S1 Table: DESeq2 and by *in situ* hybridisation (Oda-Ishii et al., 2016)). An enrichment of an Achaete-scute like binding sites was specifically observed for A-line neural genes (AN>AM). The CAG half-site is recognised as a DNA binding target by a subfamily of the class ‘A’ bHLH transcription factors (De Martin et al., 2021). The genome-wide survey of the *Ciona* bHLH transcription factors has revealed a handful of genes, whose human orthologues exhibit a preferential binding to the CAG half site. These include *Ascl.a* (KH.L9.13), *Ascl.b* (KH.C2.880), *Ascl.c* (KH.C2.560), *Atoh* (KH.C8.175), *Atoh8* (KH.C9.872), *Hand* (KH.C14.604), *Hand-r* (KH.C1.1116), *Mrf* (KH.C14.307), *Ptf1a* (KH.C3.967), *Ptf1a-r* (KH.L116.39), and *Tcf3* (KH.C3.348) (De Martin et al., 2021; Satou et al., 2003). Inspection of our scRNA-seq dataset revealed appreciable read counts only for *Tcf3*, which are observed in all the cell types analysed (see supplementary link), indicating its ubiquitous expression. Its functional involvement in segregation of the neural lineages remains to be addressed. Zic-r.b (ZicL)-like motifs were enriched in gene sets from all comparisons. *Zic-r.b* is present as multi-copy genes in the *Ciona* genome (Satou and Imai, 2018). Enrichment of ZicL-binding sites in neural lineages is consistent with the observation that *Zic-r.b* transcripts are detected in our DESeq2 analyses in both neural lineages at the neural plate stage (aN>aE and AN>AM) (S1 Table: DESeq2; Fig 4) and that *Zic-r.b* is necessary for the expression of *CELF3/5/6* (KH.C6.128) (Imai et al., 2002). *Zic-r.b* also plays a critical role in the acquisition of notochord fates (Imai et al., 2002), which is consistent with the enrichment of its binding site in AM-specific genes and its high read counts (KH.S816.1, KH.S816.2, KH.S816.4, KH.L59.1, and KH.L59.12) in the notochord lineages (A7.3, A8.5, and AM) (see supplementary link; Fig 4) (Winkley et al., 2021). It is puzzling however that the binding site is also enriched in the epidermal lineage, in which read counts for *Zic-r.b* transcripts are negligible throughout the stages analysed in this study (see supplementary link; Fig 4). Finally, binding sites of a variety of ETS-domain transcription factors were found enriched in a-line neural genes (aN>aE) and in those in the mesoderm (notochord) lineage (AM>AN) (Winkley et al., 2021). This is consistent with acquisition of these fates requiring ERK signals (Hudson et al., 2003; Yasuo and Hudson, 2007), which, as described above, can control ETS-domain transcription factors by direct phosphorylation (Yang et al., 1996).

**Fig 6.**
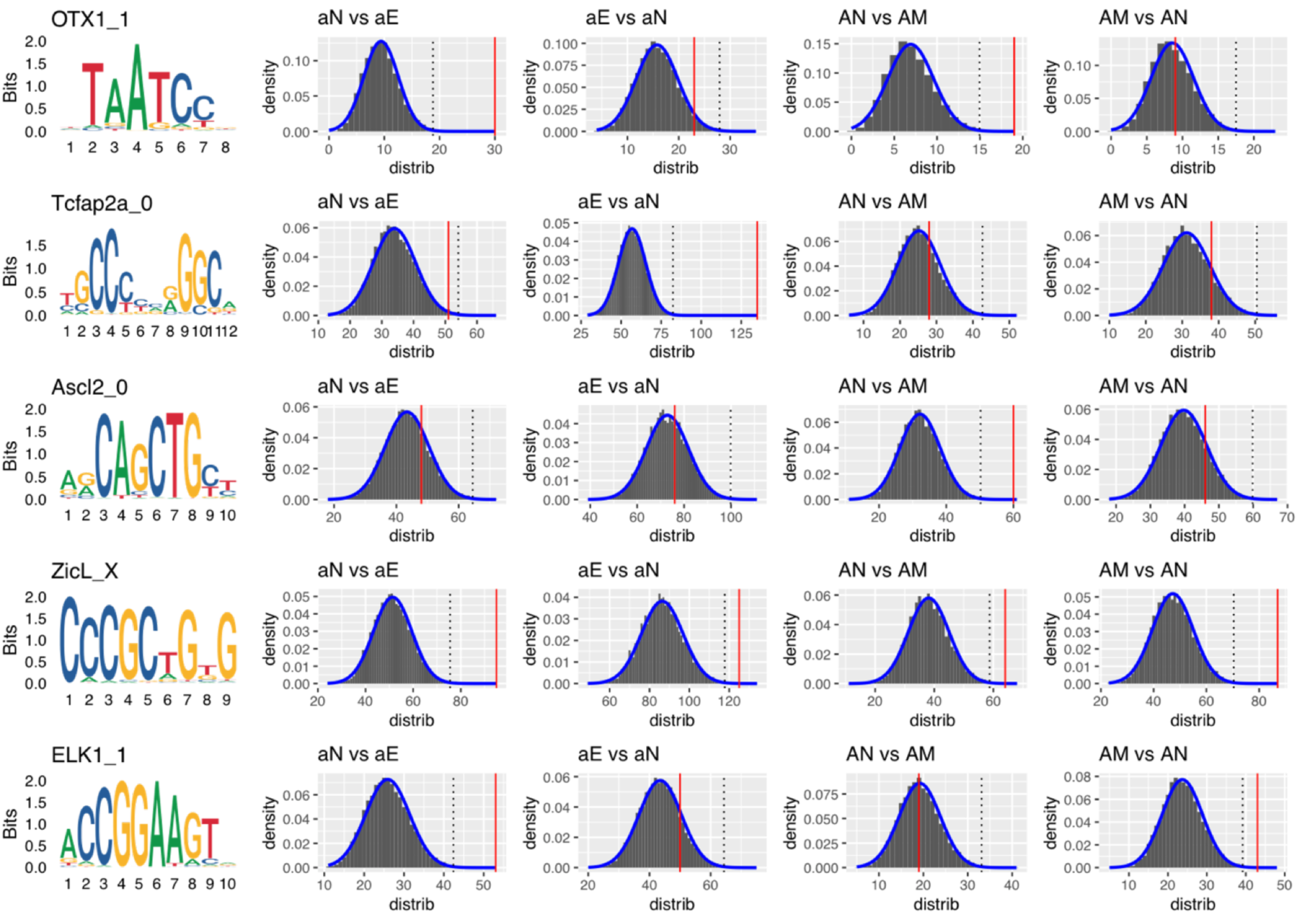
Transcription factor-binding site identification and enrichment analyses indicate distinct regulatory inputs to activate neural plate stage genes between the two neural lineages. Binding motif sequence logos are on the left column with names of corresponding transcription factors. The red line in each histogram indicates the observed number of TF sites in the 1kb upstream region of the set of significantly ‘up’ lists of genes. Numbers of genes are analysed: 131 for aN vs aE; 220 for aE vs aN; 98 for AN vs AM; 120 for AM vs AN (e.g., “aN vs aE” indicates a set of genes up-regulated in aN compared to aE). The distributions show the total number of sites from the same number of genes drawn randomly (10,000 trials) from the *Ciona* gene set. Dotted lines represent 3 standard deviations above the mean of each fitted normal distribution (blue lines). See methods for more details.

Overall, at this stage of development, the TFBS enrichment is unique for each lineage, including for the two neural lineages, consistent with the distinct ontologies of the two neural lineages. However, we observed that binding sites for both Zic.r-b and Otx are enriched in differentially expressed genes in both neural lineages.

## Conclusion

In this study, we show that neural cells become transcriptionally more similar to each other than they are to their early segregating sister lineages by the neural plate stage of development (Fig 2). We identify a set of both known and novel genes which are enriched in the two neural lineages (Fig 4). Among the shared set of genes are several which are also reported to be expressed in the CNS of vertebrates, such as *Noggin*, *ZCCHC24*, *miR-124*, *Celf*, *Zic* and possibly *miR-219,* suggesting an evolutionary important gene set (Kang et al., 2012; Knecht et al., 1995; McMahon et al., 1998; Vidyanand et al., 2017; Wu et al., 2019). In terms of regulation, Zic-r.b and Otx binding sites are enriched in differentially expressed genes of both neural lineages, suggesting an early generic requirement for these factors in neural fates. Otherwise the enrichment of transcription factor binding sites is unique in each lineage consistent with their different ontologies and specification mechanisms. We suspect that our list of shared enriched genes may be an underestimate since the number of cells analysed in any given comparison was small and read counts often showed high variance across cells, e.g. the case of *Otx* (S1 Fig). The identification of known markers, however, such as *CELF3/5/6*, *Zic-r.b* and *Pans/miR-124* indicates that the strategy used is justifiable.

## Materials and Methods

### Ascidian embryo culture and basic methods

We used *Ciona robusta* for dissection and sequencing analysis. General culture and methods of ascidians are published in (Sardet et al., 2011). Embryos were dissected on a 1.5% agarose-coated dish with a fine glass ‘hair’ smeared with cellular debris. A single isolated cell was transferred in 0.5µl of seawater into a 0.2ml PCR tube containing 9µl RNase-free water. After visual inspection of the presence of the cell under stereomicroscope, 1µl of 10x Reaction buffer (SMARTer Ultra Low Input RNA Kit, Takara Bio) was added and the tube was vortexed immediately and stored at –80°C before being processed for library preparation. For all *in situ* hybridisation (ISH) analysis, *Ciona intestinalis* was used following the published protocol (Hudson, 2020). DIG-labelled probes were made from the following cDNA clones: GC32e02 (KH.C7.140); GC11m08 (KH.C14.33); GC31n24 (KH14.310); GC03h09 (KH.C4.90); GC27k22 (KH.C12.562) (Satou et al., 2002). The probe for KH.L18.113 was synthesised from PCR fragments amplified from VES105_E20 (Gilchrist et al., 2015) using the following PCR primers: KH.L18.113-F: ATACAAGCAACTCAACCAACGC; KH.L18.113-R: *CTCACTATAGGG*TATCTTGGTCGTTCGTTTCGTC; T7-probe: ggccTAATACGACTCACTATAGGG. For *miR-124* detection, specific locked nucleic acid (LNA) probes doubly-labelled with digoxigenin were purchased from Qiagen and ISH was conducted following the published protocol (Chen et al., 2011). ISH was not attempted for KH.C11.540, KH.C1.29 or KH.C7.59.

### Library preparation and sequencing

Indexed Illumina libraries were prepared using Nextera XT kits (Illumina) from cDNAs generated with SMARTer Ultra low input RNA kit (Takara Bio) according to manufacturers’ instructions. Sequencing was conducted in the UCA GenomiX genomics platform (Sophia Antipolis, France) using Illumina NextSeq500 system. For each library, single end 75bp reads were obtained with sequencing depth of around 8 million reads par cell.

### Computational analysis

*Ciona robusta* sequences and annotation were taken from the GHOST database (http://ghost.zool.kyoto-u.ac.jp/download_kh.html) – specific files used are named below.

#### (1) Read mapping & counts

Reads were aligned to the *Ciona* genome using STAR (version 2.7.10a) with the ‘--quantMode GeneCounts’ option to quantify reads per gene (Dobin et al., 2013). Exon annotation was taken from the KH.KH.Gene2013.gff3 file reprocessed for compatibility with STAR gene/exon format. ReadsPerGene files were merged to a single gene by cell type raw count matrix.

The count matrix of raw reads is available at http://github.com/rcply/ciona_sc/

#### (2) Differential expression analysis

The raw count matrix was analysed using DESeq2 (Love et al., 2014). Two outlier cells were consistent outliers and excluded from further analysis (A61l5 and A87r5). Comparisons were performed using count matrices restricted to the particular stage (e.g. only cells from the 32-cell stage). To deal with batch and animal specific effects reflecting data collection constraints, comparisons involving the 16-, 32-and 64-cell stages included an animal of origin term in their design formula. DESeq2 alpha was 0.01 and log fold change cutoff 0.0. lfcShrink was performed using ‘ashr’.

The PCA plot data were first subjected to Variance Stabilizing Transformation (using the ‘vst’ function) followed by the plotPCA function of the DESeq2 package, using the default top 500 genes by row variance.

The cluster tree of cell types was produced from within the Seurat package (Stuart et al., 2019), using the BuildClusterTree function with a PCA reduction (from Seurat) with dimensions 1 to 12.

The code for these tests is accessible at https://github.com/rcply/ciona_sc/

#### (3) miR-124 target site enrichment

3’UTR exons were taken from the KH_Ciona_2013.fa file using annotation in the KH.KHGene2013.gff3 file. Transcripts were processed individually. For each transcript exons labelled as ‘three_prime_UTR’ were concatenated and the complete UTR sequence reverse complemented where necessary. These sequences were searched for the ‘7mer-m8’ (GTGCCTTn) and ‘7mer-A1’ (nTGCCTTA) *mir-124* target sequences (which together also cover the ‘8mer’ sequence), taken from (Chen et al., 2011). Sequence identifiers were sorted for redundancy at the locus (i.e. gene) level. This resulted in a list of 873 potential *mir-124* targets. Chi-square tests were performed on the intersection of this list with the 4 neural plate stage cell-type differential expression lists.

#### (4) Transcription factor binding site enrichment

We searched regions of the *Ciona* genome 1kb upstream of gene start sites based on the KH.KHGene2013.gff3 annotation, after first filtering to remove low complexity regions as defined by nseg (Wootton and Federhen, 1993) (https://github.com/jebrosen/nseg). Transcription factor binding site models (n=745) were taken from the profile weight matrix (PWM) data described in (Jolma et al., 2013). Although these data are not derived from *Ciona*, transcription factor specificity is generally well conserved over longer evolutionary timescales (Nitta et al., 2015). Because of its recognised importance in early neural differentiation, we manually added the *Ciona* ZicL (Zic-r.b) binding motif taken from data in Figure 2 of (Yagi et al., 2004). PWM searches were performed using the MOODS software library (Korhonen et al., 2017), with a false positive rate of 0.0001. Exactly overlapping matches to the same PWM on forward and reverse strands (i.e. palindromic sites) were counted as one match.

For each pairwise cell comparison, a set of ‘real’ genes was taken as those with significant DESeq2 p_adj values. For each PWM, the number of binding sites in the ‘real’ set and the number of binding sites in the remaining set were compared using a nested Poisson model to determine if they occurred at a significantly different rate in the ‘real’, with a P-value adjustment for testing multiple PWMs. For PWMs with an adjusted P-value more significant than 0.05, we also generated 100 permuted ‘real’ sets and excluded PWMs where the P value from the ‘real’ set was ever found to be less significant than any permuted value, to exclude cases where this model is inappropriate – these sites are marked FALSE in S3 Table.

For visualization purposes, we generated plots of the total number of binding sites for a particular PWM from 10,000 randomly sampled sets of genes of the same size as the DESeq2 ‘real’ set for a particular comparison. We fit a normal distribution to these plots and mark the true number of binding sites for the real set.

The code for these tests is accessible at https://github.com/rcply/ciona_sc/.

#### (5) mir-124 target site enrichment

3’UTR exons were taken from the KH_Ciona_2013.fa file using annotation in the KH.KHGene2013.gff3 file. Transcripts were processed individually. For each transcript exons labelled as ‘three_prime_UTR’ were concatenated and the complete UTR sequence reverse complemented where necessary. These sequences were searched for the ‘7mer-m8’ (GTGCCTTn) and ‘7mer-A1’ (nTGCCTTA) *mir-124* target sequences (which together also cover the ‘8mer’ sequence), taken from (Chen et al., 2011). Sequence identifiers were sorted for redundancy at the locus (i.e. gene) level. This resulted in a list of 873 potential *mir-124* targets. Chi-square tests were performed on the intersection of this list with the 4 neural plate stage cell-type differential expression lists.

## Acknowledgements

We thank Nori Satoh, Ute Rothbacher and colleagues for the *Ciona* Gene Collection plates. We thank Bob Zeller for help and advice with the LNA-miR-124 ISH protocol and Chen Cao for comments on the manuscript. The project was funded by an Emergence program of Sorbonne Université (SU-16-R-EMR-67), which also supported JB. The teams of HY and RRC were supported by the Centre National de la Recherche Scientifique (CNRS), Sorbonne Université, the Fondation ARC pour la Recherche sur le Cancer (PJA 20131200223 to HY) and the Agence Nationale de la Recherche (ANR-17-CE13-0003-01 to HY). All image acquisitions were conducted in the imaging platform PIM (member of MICA). Our imaging platform (PIM) and animal facility (CRB) are supported by EMBRC-France, whose French state funds are managed by the ANR within the Investments of the Future program under reference ANR-10-INBS-0.

## Supporting information

**S1 Fig.**
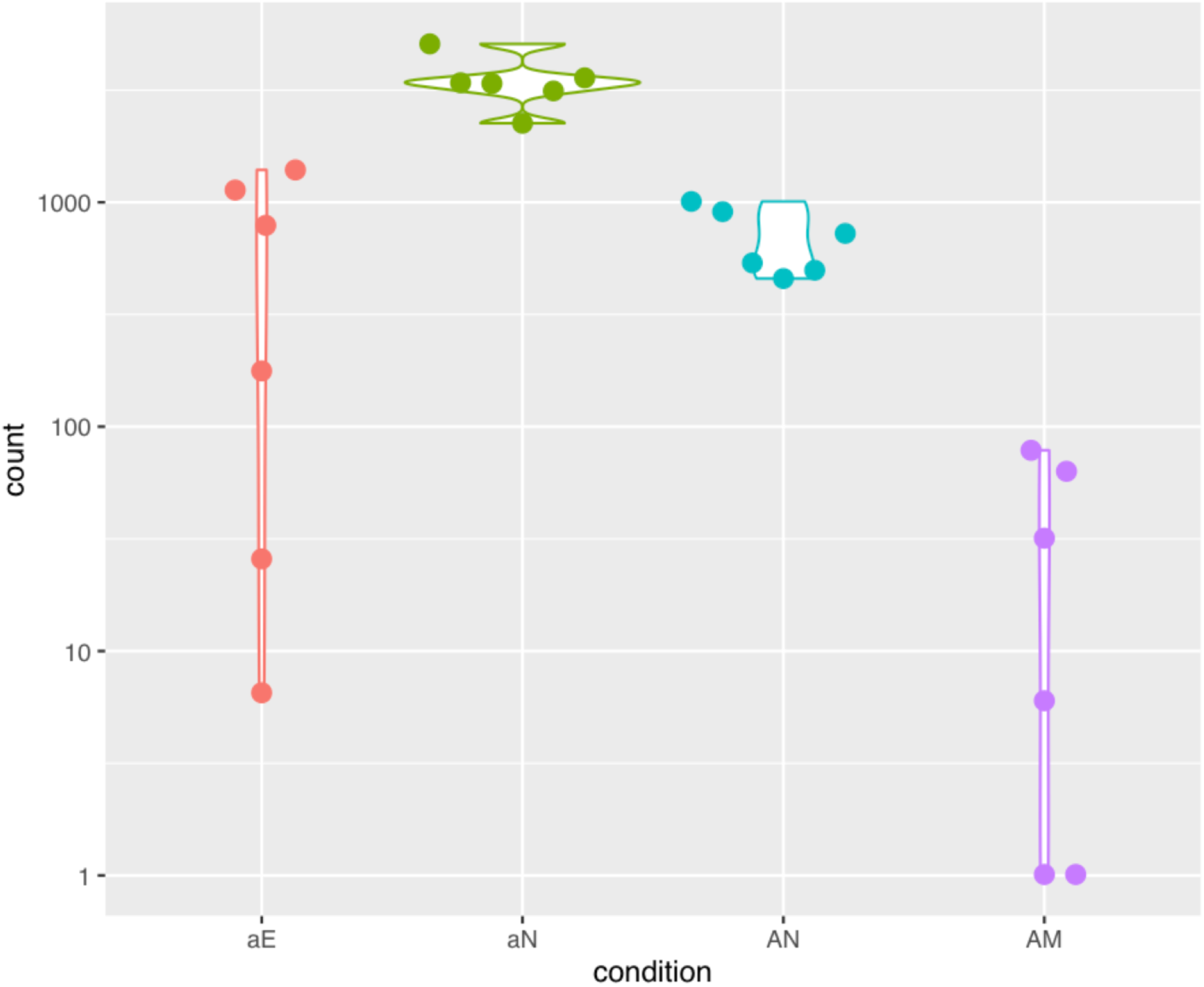
Expression of *Otx* at neural plate stage. Although there is clear separation between the expression levels in AN and AM (right hand side, blue, purple), this is less obvious in aN vs aE (left hand side, green, red) owing to the high variability of aE counts (note log scale).

**S1 Table. DESeq2 results**.

**S2 Table. Shared neural genes.**

**S3 Table. Enriched transcription factor binding sites.**

## Notes

https://github.com/rcply/ciona_sc/

### Competing Interest Statement

The authors have declared no competing interest.

